# Longhorn Beetles Form Structural Colour Using Calcium Phosphate Biominerals

**DOI:** 10.1101/2024.10.01.616160

**Authors:** Yin Chang, Hsiang-Han Tseng, Masahiko Tanahashi, Darius Pohl, Bernd Rellinghaus, Luca Bertinetti, Yael Politi

## Abstract

Brilliant structural colors originating from diverse photonic crystals are found across many phyla, including the striking iridescent colors of beetles and butterflies, produced by three-dimensional photonic crystal structures in the specialized cuticular scales. However, the precise composition of these structures remains largely unknown, although it is key to unravelling colour production mechanisms and morphogenesis. The longhorn beetle *Doliops similis* displays vibrant green patterns on its otherwise dark elytra. These patterns are formed by arrays of minute scales that encompass a three-dimensional photonic crystal made of orderly packed nanospheres. We found that these nanospheres are composed of carbonated amorphous calcium phosphate biomineral. By accurately parameterizing the structure and calculating the refractive index deduced from the relative fractions of the organic and inorganic phases, we derived reflection wavelengths that match the observed green hue, demonstrating the biomineral’s role in colour production. Our study further reveals that biomineralization is widespread in the Lamiinae subfamily, with colour diversity achieved through variations in nanosphere size, packing, and composition. This study opens new avenues for developing bioinspired mineral-based optical devices with high refractive indices and defect-resistance, overcoming the shortcomings of current polymer-based designs.

## Introduction

Many organisms synthesize minerals utilizing ions imbibed from the environment by absorbing ions from the environment and depositing them in specialized cellular compartments^1,2^. Biomineralization is widespread across bacteria, plants, and animals^3-5^. Within the arthropod phylum, biomineralization is widespread within marine and terrestrial crustaceans. These organisms use hierarchically organized chitin fibres that are cross-linked with proteins and heavily mineralized with amorphous calcium carbonate (ACC) ^7-10^, amorphous calcium phosphate (ACP)^1^, and, in some cases, small amounts of crystalline minerals like calcite and carbonated hydroxyapatite^6^. However, among insects, biomineralization is rare^8,9^ and primarily observed in the cuticles of larval and pupal dipterans^11-14^, some tenebrionid beetles^15^, and fungus-farming ants^16^.

Although some biomineral arrays, such as mollusk nacre, also known as mother of pearl, show structural coloration, functional biomineralised photonic crystals are uncommon, with the blue-rayed limpet (*Patella pellucida*) ^17^ being one of few known examples. More commonly, organisms use organic-based photonic crystals^18-29^, including for example, the guanine arrays in fish, geckos and spiders^18^, laminar or crystal-like keratin and melanosomes in bird barbules^19^, multi-layered cuticles in insect exoskeletons^21^, bi-continuous periodic lattices in the scales of weevils and butterflies^23-29^ and more.

Here we show that amorphous carbonated calcium phosphate (hereafter termed ACCP) nanospheres are the main component of the photonic crystals in the scales of the longhorn beetle *Doliops similis*, responsible for its metallic green colour due to the nanospheres’ uniform size, ordered packing, and high refractive index. Additionally, biomineral photonic crystals were found to be widespread within the Lamiinae subfamily, displaying a variety of colours. These findings highlight previously under-appreciated role of biomineralization in insect coloration and demonstrating the diversity of bio-photonic crystals in nature.

### Brilliant green structural colour from highly ordered opal-structured photonic crystals

The brilliant green patterns of *Doliops similis* are produced by micron-sized scales, connected to the black exoskeleton at one end by a narrow stalk (**SI Fig. S1**). The scales generate angle-dependent green colour, which gets slightly blue-shifted at high specular angle, giving an iridescent effect (**Fig. 1a**). A single scale removed from the black elytra and placed on glass (**Fig. 1b**) shows green reflection centered around 520-550 nm. Unlike pigmented colours, structural colours arise from light interacting with periodic dielectric structures, as dictated by both their lattice and composition. Scanning electron micrographs (SEM) of a broken scale reveals its core is filled with tightly-packed nanospheres (**Fig. 1c**). Backscattered electron micrographs reveal that native photonic crystals (PC) emit a stronger signal compared to the skin, cuticles, or resin, indicating their distinct composition, comprising a material of higher average atomic number relative to organic matter. This difference disappears after heavy metal staining (**Fig. 1d**).

**Fig. 1:**
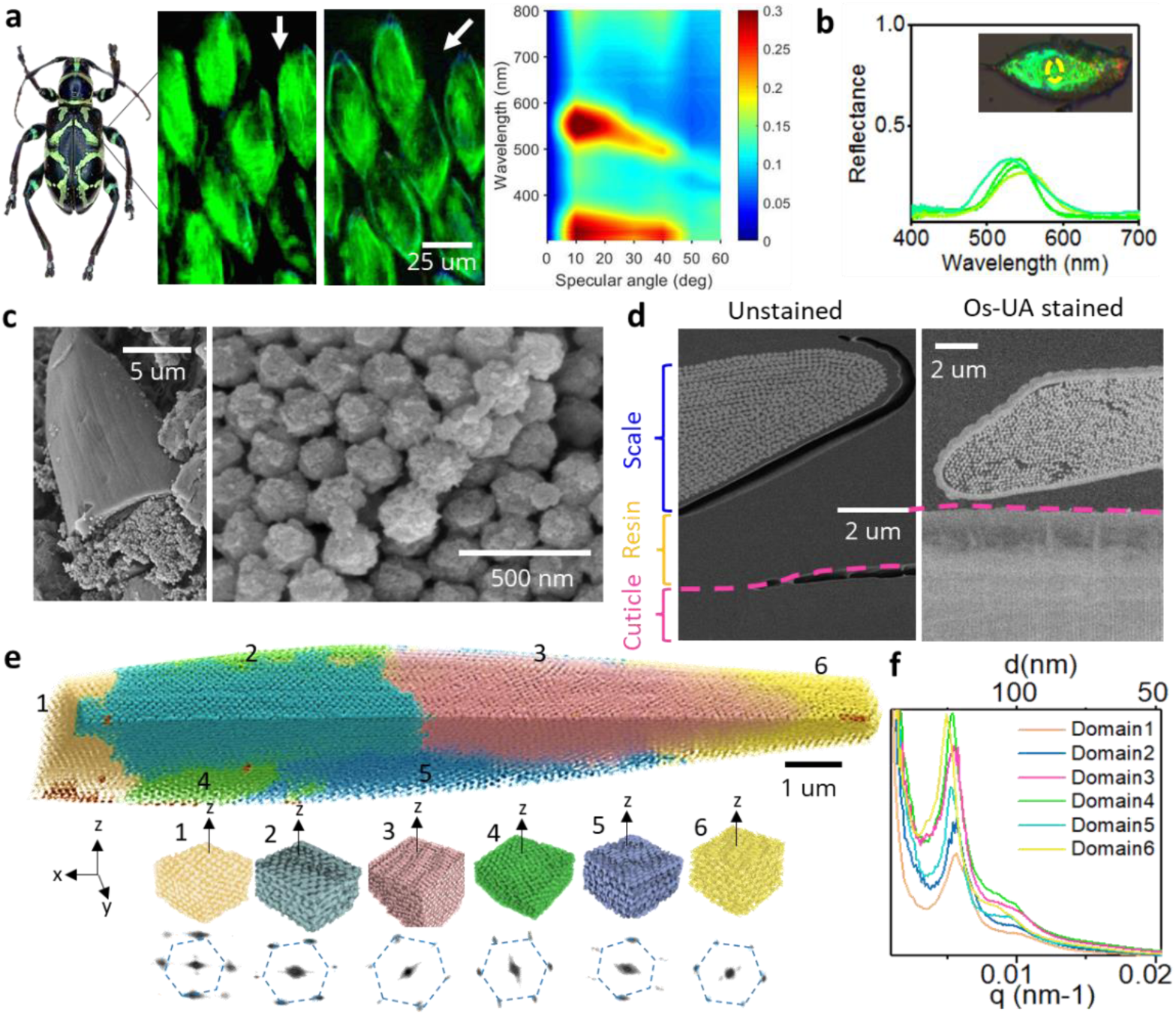
Structural colour and photonic crystals (PC) in the scales of *Doliops similis*. **a**, The adult beetle (left), its scales under varying illumination angles (arrows, middle) and specular spectra (right) measured at different angles (5 mm spot size). **b**, Reflection spectra from a single scale on a glass substrate measured using an optical fibre (few *μ*m spot size, yellow circle). **c**, SEM images of a broken scale and the inner core PC. **d**, Backscattered electron SEM images of a cross-section of resin embedded scales and cuticles. Left: native, unstained sample. Right: sample stained with heavy metal ions (Os: Osmium tetroxides; UA: Uranyl acetate). **e**, Top: Volume rendering of the PC segmented from a FIB/SEM image stack of a single scale. Different colours indicate coherent crystallographic domains. Bottom: Representative small volumes of each domain reoriented with the normal to their most densely packed plane parallel to the vertical z axis. Below: a virtual plane (normal to the z axis) through the 3D FFT of each domain. **f**, polar-azimuthal integration of the 3D FFT of each domain (the color code of spectrum is the same as corresponding domains).

High-resolution volumetric analysis (isotropic voxel size of 10 nm) of the inner core of an intact scale reveals that the nanospheres are orderly packed, forming an organized lattice comprising several subdomains. Analysis of the 3D Fast Fourier Transform (FFT) of coherently oriented domains (**Fig. 1e)** indicates a similar symmetry for all domains (**Fig. 1f)**. By stacking micrographs of nanospheres within sequential maximum density planes, we identify that adjacent regions, even within the same domain, show face-centered cubic (FCC) and hexagonal close-packed (HCP) symmetries (**SI Fig. S2**). Apart from its two tips, within one scale, we typically find two distinct domains between the upper and lower skins, meeting roughly at its centre. Domains attached to the upper skin (numbered 2, 3, 4, 6 in **Fig. 1e**) display near-perfect hexagonal symmetry, with their planes of maximum density (i.e., (111) for FCC or (001) for HCP) being parallel to the scale surface. In contrast, domains 1 and 5 show distorted hexagonal patterns (**Fig. 1e**).

The filling fraction (FF) of most domains are in a range of 63 to 67 % apart from domain 1 with FF of 55%. Using the FF and the d-spacing of (111) plane (d_111_) derived from integrated FFT spectra, we calculated diameter of nanospheres (∅) and lattice constant (*a*) of individual domain (**SI Table S1**, readers are referred to the **SI section 1-2** for details). We observed a correlation between the structural features and the positions of domains within a scale; Smaller nanosphere and lower d_111_-spacing are observed in domains closer to the stalk as well as in domains attached to the upper side of a scale (1, 2, 3), relative to those at the lower side (4, 5, 6). Despite these variations, the nanosphere diameter was homogeneous (∅: 218± 35.78 nm), and the lattice spacing, d_111_= 187.9± 26.5 nm along the scale surface normal, is uniform over the whole scale. A notable exception is seen in domain 6 where nanospheres size and shape as well as d-spacing varied substantially, especially closer to the tip of a scale (**SI Fig. S3**) with larger sphere diameter and lattice constant.

### The core of the scales is comprised of a biomineral

Intrigued by the high BSE contrast of the PC, we further analysed the compositions of the PC and the skins of scales using depth-profiling X-ray Photoelectron Spectroscopy (XPS) and Time of Flight Secondary Ion Mass Spectrometry (ToF-SIMS), by homogeneously removing biomaterials with Ar^+^ cluster etching (**Fig. 2a**, see methods). Variations of abundant atoms, with tens of nanometres depth resolution, are determined from XPS depth-profile (**Fig. 2b**). The dominating C, N, O (**Fig. 2c**) are expected as the most abundant elements in chitin and proteins typically forming the cuticle. The different relative fractions of these elements between the skin and the cuticle, suggest that the composition of the skin is somewhat different. In the PC, O is the most abundant element (40%), followed by Ca, C and P.

**Fig. 2:**
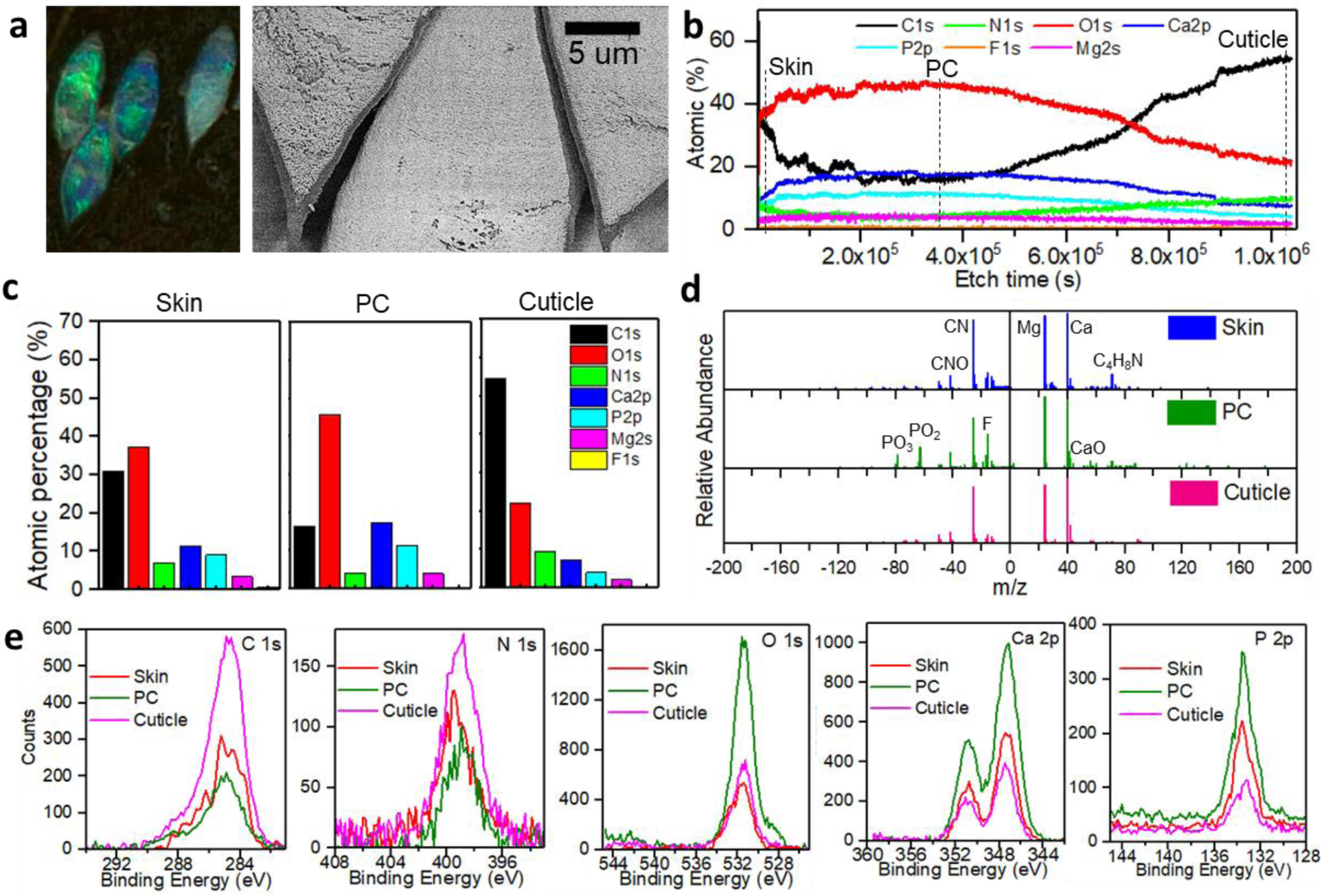
Depth profile analyses of compositions of skin of scales, PC, and cuticle. **a**, Post-Ar^+^ cluster etching sample reveals persistent structural colour (left) and well-preserved PC structures (right, BSE image). **b**, Depth profile of elemental composition across the scales and the cuticle, derived from XPS peak integration. Note: The profile shows the variation of element content with continuous etching time, which is not proportional to the thickness of the samples. Three etch times (dashed lines) were selected to represent the transition between the scale and the upper skin, the PC, and cuticles for generating **c**, the histogram of elements; **d**, ToF-SIMS depth profile: Mass spectra of negatively charged compositions (left panels with negative m/z) and positively charged compositions (right panels with positive m/z). Note: The intensity of m/z across different channels is not quantitatively comparable; **e**, XPS spectra of C 1s, O 1s, Ca 2p, P 2p scans.

The high yields of PO_2_ ^-^ and PO_3_ ^-^ as well as Ca and Mg within the PC detected by ToF-SIMS (**Fig. 2d**), strongly point towards a calcium phosphate mineral. CN^-^ and CNO^-^, possibly from proteins and/or chitin, are detected in all regions of a scale. Over whole scales, very low counts for m/z > 100 imply the scarcity of long-chain lipid or fatty acids, which normally show periodic fingerprints from m/z>250^30,31^

The binding energy (BE) of each element contains further information on the molecular species in each region (**Fig. 2e**). The broad C 1s spectra indicate the presence of various carbon species. In comparison to the cuticles, lower signature of aliphatic carbon (BE at 285 eV) is detected in the skin and PC. Carboxylic acid or carbonate (O-C=O, BE ≈ 289 eV) are not detected in the skin. Thus, not only the amount but also the type of organic material seems to vary between the skin and the core of the scale and both differ from the beetle elytra cuticle. The presence of mineralized calcium phosphates and carbonates are supported by the BE of Ca, Mg, P and O signals. Deconvoluting the O 1s spectra over the depth profile (**SI Fig. S4a, b**) reveals that in addition to these organic and inorganic oxygen species (BE ≈ 531-532 eV), small amount of hydroxides (5 %, BE at 533 eV) and metal oxides (MgO and CaO, <5 at%, BE at 529 eV, **SI Fig. S4c**; **Fig. 2d**) are present in the skin and PC of scales. These alkali-earth metal oxides are not naturally stable and are likely decomposition products of their hydroxides due to the water loss under ultrahigh vacuum at room temperature^32^. The Ca 2p and P 2p spectra as well as the calculated Ca/P ratio of 1.5 and O/P ratio of 4.4 in PC are close to the theoretical values in amorphous calcium phosphate [Ca_x_H_y_(PO_4_)_z_·nH_2_O, H_2_O≈20wt%, and Ca/P ratio 1.2–2.2, ACP] or α-tricalcium phosphate [α-Ca_3_(PO_4_)_2_, α-TCP] (**SI Table S2**). CaCO_3_ typically show strong satellite loss in Ca 2p at 355 eV and 359 eV, which were not observed in scales (**SI Fig. S5**).

### The photonic crystal is made of high refractive index amorphous calcium phosphate mineral

Using X-ray diffraction (XRD) and Fourier-transform Infrared Spectroscopy (FTIR) spectroscopy, we determined that amorphous, rather than a crystalline, carbonated calcium phosphate is the main mineral phase comprising the photonic crystals. Indeed, the 2D XRD detector images of the scales show an isotropic intensity distribution that is also independent of the sample rotation (**SI Fig. S6**). The integrated XRD profile of intact scales shows contributions from both the skin and PC (**Fig. 3b**), with the PC profile displaying amorphous rings and two broad peaks at q=21 and 32 nm^−1^, comparable to synthetic ACP. The skin profile exhibited peaks at q=6.3 and 13.7 nm^−1^, typical of proteinaceous materials^33,34^, and it differs from the cuticle, which show clear chitin reflections and protein background^35^. Furthermore, the strong Small Angle X-ray Scattering (SAXS) signals from PC and ACP result from their nanospheres morphology and the high electron density contrast against air filled voids.

**Fig. 3:**
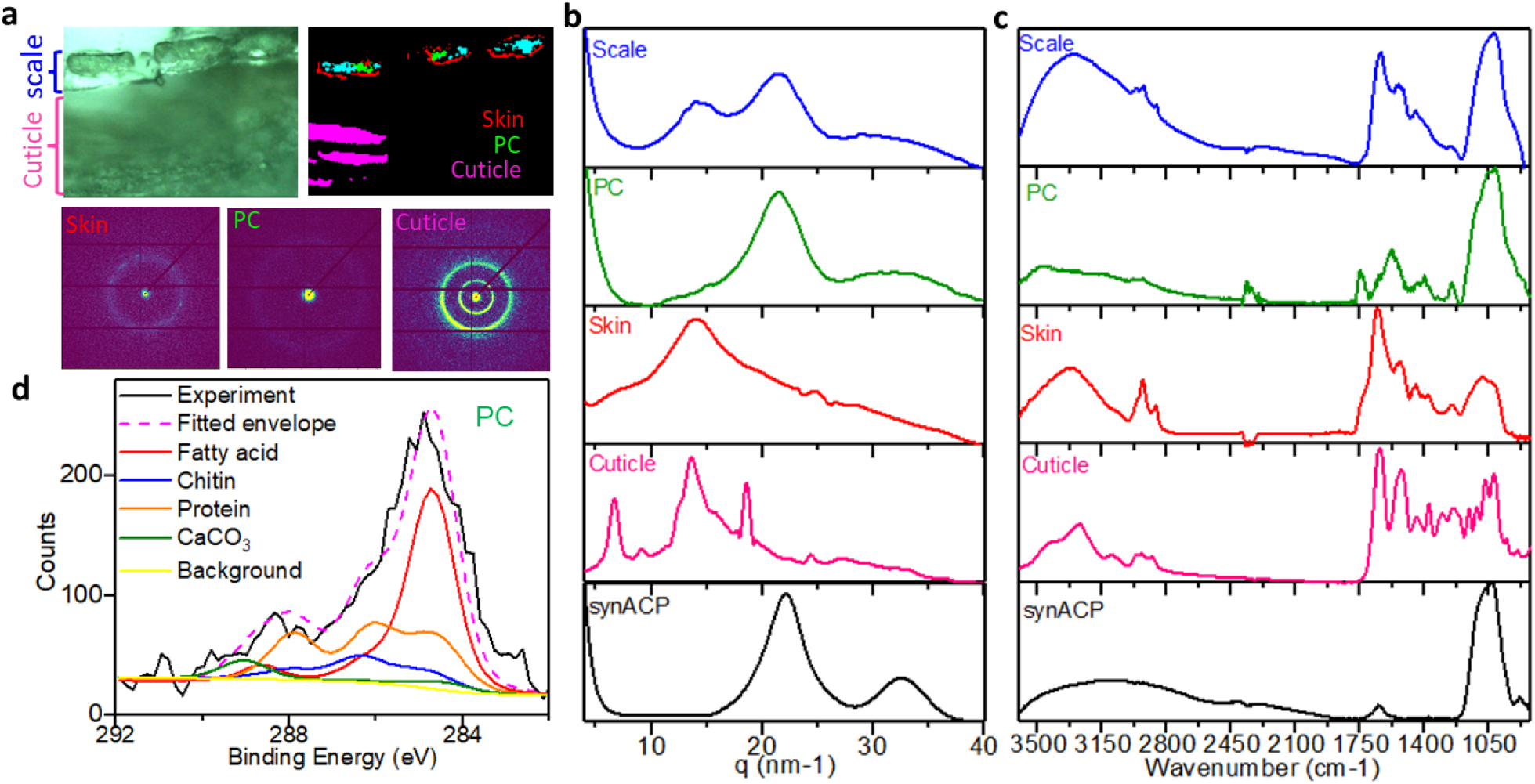
Materials characterization of intact scales, scale skin, PC, and cuticles by XRD, FTIR, and XPS multivariate analysis. **a**, The sample cross-section for SAXS/WAXS measurement^50^ and masks produced by difference of diffraction patterns. **b**, Integration of XRD signals of skin (red mask), PC (green mask), and cuticles (magenta mask) over a cross-section of scales attached to the cuticles. **c**, The FTIR spectra of intact scales (blue), isolated PC (green), isolated scale skin (red), cross-section of cuticles (magenta), and synthetic amorphous calcium phosphate (synACP, black). All XRD and FTIR spectra were normalized. **d**, Non-linear least squares fitting (magenta dash line) of the experimental XPS C1s spectra (black line) of PC with multivariate components of fatty acid, chitin, protein, and CaCO_3_.

FTIR spectra of the different organic parts of the scales show the typical signature of the common building blocks of insect’s cuticle: fatty acids, chitin and protein. Characteristic amide I and II peaks (respectively around 1650 cm^−1^ and 1550 cm^−1^), typical of protein/chitin, are observed in both the skin and elytron cuticle, but the skin shows a higher protein ratio and more fatty acids, indicated by a pronounced shoulder at 1750 cm^−1^ and a strong peak at 2900 cm^−1^. Instead, the spectrum of the PC shows mainly strong phosphate peaks, associated to HPO_4_ ^3-^ and PO_4_ ^3-^ at 850-1200 cm^−1^. The shape of this band resembles very closely the one of synthetic ACP (**Fig. 3c**), in agreement with the XRD profile. Although chitin also has contribution in this range, the PC spectrum lacks other chitin contributions, specifically in the amide regions; instead, two peaks at 1750 and 1250 cm^−1^ are observed, which can be assigned to the stretching modes of C=O and C-OH of the protonated carboxyl group most likely associated to fatty acids^36^. Characteristic peaks of CO_3_ ^2-^ stretching are weak but observable at 1071 cm^−1^ and between 1400 and 1480 cm^−1^, implying the presence of small amounts of carbonate within the mineral phase. In addition, the broad water bending peak centred at around 1630 cm^−1^ suggests that the mineral is hydrated.

By means of non-linear least squares fitting of the XPS C 1s spectra using reference multivariate^37^ (**SI Fig. S7**), we could further attribute the 16.5 at% carbon in PC (**Fig. 3d**) to fatty acids (7.8 at%), chitin (2.1 at%), protein (5.6 at%), and CaCO_3_ (1 at%). From these fractions and the theoretical ratio of N, O, and C in different biomolecules (**SI Table S3**), the relative contribution of organics, calcium carbonate, and calcium phosphate phases in the PC are estimated to be about 11%, 9%, and 80% (see calculations in **SI section 2**).

Strikingly, the nanospheres themselves are also not homogeneous. Robust Principal Component Analysis (RPCA) of the Transmission Electron microscopy-Electron Energy Loss (TEM-EELS) spectra in the range 220-500 eV revealed that the core of the ACCP nanosphere (**Fig. 4** red) contains increased carbonate contribution (sharp *π*\* peak at 290 eV, and *σ*\* at 301 eV), whereas the shell (**Fig. 4** green) and some pixels covering the sphere surface (**Fig. 4** green), shows shoulder peaks at 287 eV, 285 eV, as well as the broad signal in the *σ*\* region, consistent with organic matter^38,39^. The core-shell structure is also reflected by elemental mapping using energy dispersive spectroscopy (EDS), showing that the carbon signal is higher in the shell as opposed to the core of the nanospheres (EDS, **Fig. 4c**, and **SI Fig. S9** for TEM-EDS mapping).

**Fig. 4:**
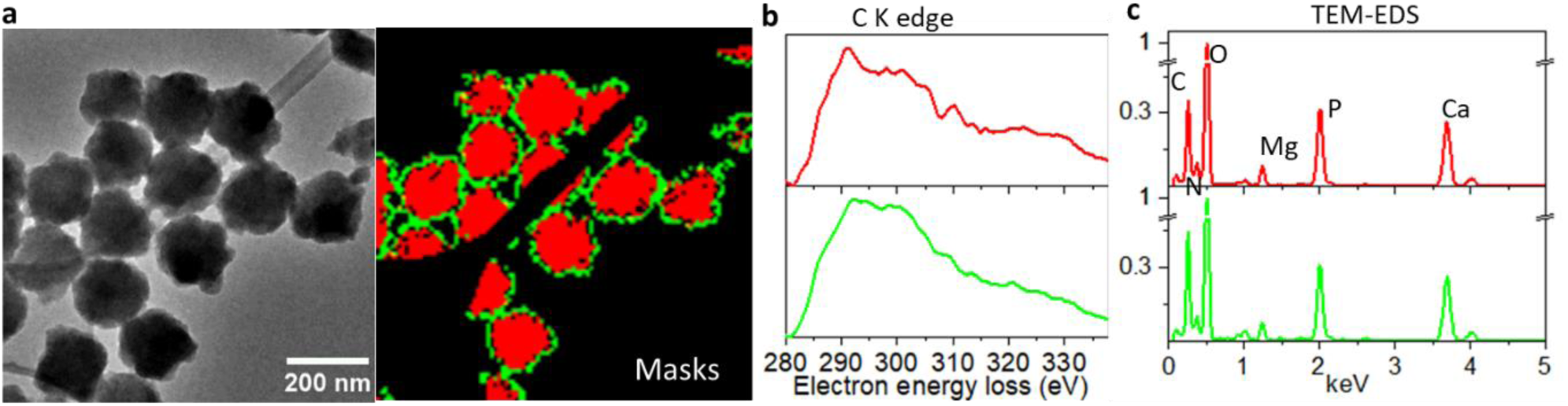
TEM energy dispersive X-ray/electron energy-loss spectroscopy (TEM-EDS/EELS) of the nanospheres. **a**, TEM image of the sample and the PCA decomposition factors (mask with different colours) of the EELS C K edge spectra. **b**, TEM-EELS C K edge spectra of each masked area (based on RPCA, see **SI**). **c**, Normalized TEM-EDS spectra from the individual mask area.

Using the composition analysis and relative component concentrations, the effective refractive index of PC is estimated as of 1.62-1.63 (for calculation details see Methods). Inserting the effective refractive index of photonic crystals and the average d_111_= 185.4± 6.3 nm and into Bragg-Snell’s law, we calculate an expected reflection peak in the range of 522.8-531.7 nm. This is in a good agreement with the visible green colour, as well as the reported specular reflection spectra (peak position at 525 nm) of a single scale of *D. similis* beetles (**Fig. 1b**).

### Biomineral photonic structures with diverse colours are common in Lamiinae

The results presented so far demonstrate that the opal-like photonic structure giving rise to vivid colouration in *D. similis* (Lamiinae: Apomecynini) is made of carbonated amorphous calcium phosphate biomineral. We found the biomineral photonic crystals are commonly produced by longhorn beetles in the Lamiinae subfamily (**SI Fig. S8**), despite large variation in structural designs (**Fig. 5**). Biominerals were observed at the tribe level, in the laminar mineral layers in Saperdini tribe as well as in isolated biomineral nanospheres embedded in fibrous organic matrices in Mesosini and Pteropliini tribes.

**Fig. 5:**
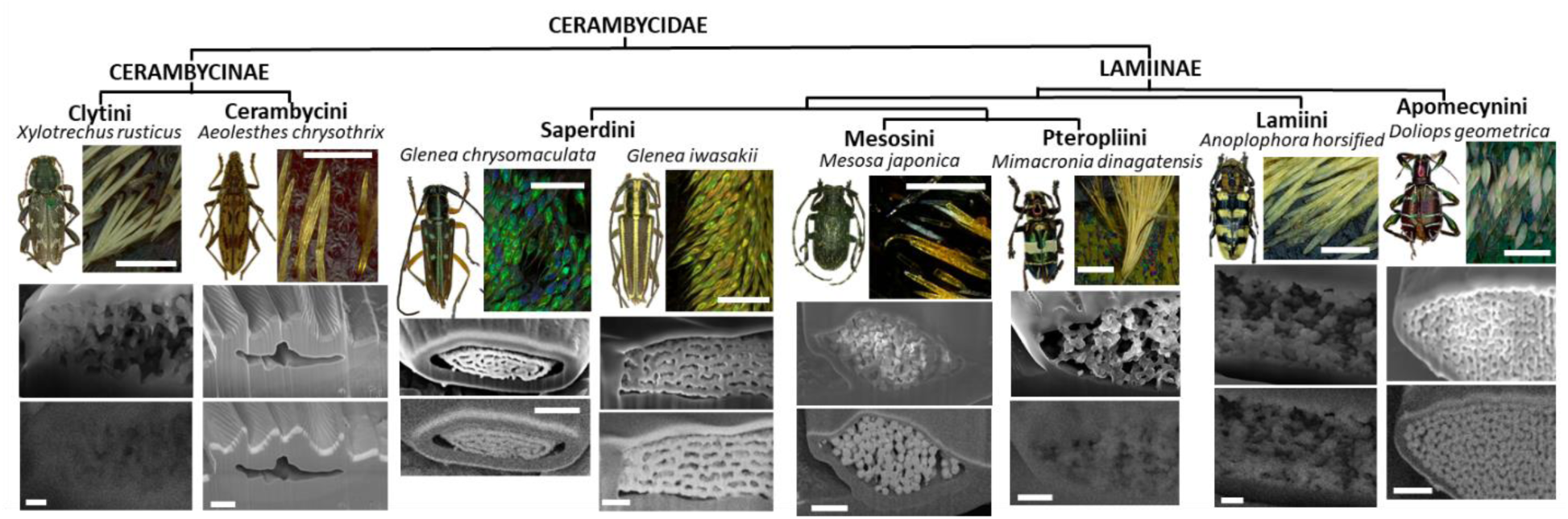
Comparison between PC in the scales of differently coloured longhorn beetles in Cerambycinae and Lamiinae subfamilies. Light micrographs of the beetles and their scales (upper row), SEM (middle) and BSE (lower row) micrographs are presented. Scalebar in light microscope images: 100 *μ*m. Scalebar in SEM and BSE micrographs: 1 *μ*m.

## Discussion

Our study indicates that *D.similis* forms three-dimensional opal-like photonic crystals with carbonated amorphous calcium phosphate mineral nanospheres. The high refractive index of ACCP minerals and the highly ordered structure including the alignment of the close packed plane (111) parallel to the scale surface are responsible for their metallic green appearance. Structural variations over domains in the PC are associated with their positions within the scale and probably affected by its shape. Each ACCP nanosphere contains a highly carbonated core and a thin organic covering which seems “gluing” neighbouring nanospheres together.

ACC/ACP biominerals in marine and terrestrial crustaceans are packed randomly and show a large size distribution (range from few hundreds of nanometres to micrometre)^7-9^. The high uniformity and order of ACCP observed in *D.similis* longhorn beetles’ scales is exceptional and raises many questions on the formation mechanisms of the photonic crystal structure: an organics pre-formed template may be involved, or particle assembly and dense parallel packing may be, in part, constrained by the encasing skin of the scale. Thus, the driving forces for the alignment may be direct nucleation, or stress due to confinement. Insects’ cuticular scales are produced by epidermal cells. The PC in butterfly scale was suggested to form templated by the cell’s smooth endoplasmic reticulum^24^. However, it is not known how the organism accumulates and secrets high Ca, Mg into the scales. Mineralization in crustacea is driven by mineral fluxes across the cuticle from various gastral storage sources. Future research on the relationships between cellular development and the formation of the scale in longhorn beetles will provide new insight on possible formation mechanisms of these structures and may reinforce the homology or dissimilarity of these mineralized scales with other cuticular scales.

Efforts have been made into engineering a bottom-up assembly of opal structures for their promising optical properties^40-42^. The low refractive index of 1.45-1.59 typical of the commonly used silica or polymeric materials, limited tunability of particle sizes, and amounts internal defects involved in the formation constrain the applications^40^. Lab-synthesized ACP/ACC nanospheres can be produced with tailored sizes, ranging from tens of nanometers up to 200 nanometers, and narrow size distribution via spinodal decomposition, sol-gel chemistry, or hydrothermal synthesis^43-45^, offering an interesting synthesis routes for PC design. Additionally, varying the amount of carbonate in ACCP solid-solutions can further influence the size of the nanoparticles. However, lab-synthesized ACP/ACC particles do not present a particular ordered assembly. It is therefore especially interesting to investigate the self-assembly of biomineralized opal-structures as observed in the *D.similis* for bioinspired PC synthesis. Further studies on the PC of differently coloured scales made by disordered minerals will also help to understand the relationship between mineral composition, particle size and the associated reflection colours.

## Acknowledgements

We thank C. Chang S. Weiner and and P. Vukusik and S. Mouchet for their helpful comments on our study. We thank T. Masahiko, T. Wataru, C-P. Lin for field work collections and drawing our attention to the intriguing features of the Lamiinae longhorn beetles. We acknowledge the European Synchrotron Radiation Facility (ESRF) for provision of synchrotron radiation facilities under proposal number 5328 and proposal number 5470 and we would like to thank J. Liu, A-A. Medjahed, M. Burghammer for invaluable assistance and support in using beamline. We thank E. Tallarek from TESCON performing ToF–SIMS measurements. We thank Y.Ogawa for preparing TEM sections and purified standards. We thank S.Vignolini for permission to use unpublished optical measurements. We acknowledge the use of the facilities in the Dresden Center for Nanoanalysis (DCN) at the Technische Universität Dresden. We thank S. Mammadov from Thermo Fisher Scientific introducing and assisting us on the XPS measurements. We thank to the support and encouragement from B. Chou, Y. Chang, M. Katze.

## Author contributions

Y.C., L.B. and Y.P. conceptualized the project and designed the experiments. Y.C. and L.B. conducted FIB/SEM measurements, reconstructed 3D volume imaging, and performed structural analyses. Y.C. and H-H.T. performed XPS measurements and analyses. D.P. and B.R. performed the TEM imaging and EDS/EELS measurements. Y.C. and L.B. performed the hyperspectral analyses of ToF-SIMS, XRD, TEM-EDS/EELS data. Y.C. H-H. T., M.T., L.B. and Y.P. wrote the original draft. All authors edited the manuscript and read and approved the final version of the manuscript.

## Data availability

All data are available in the main text, the Supplementary Information

## Materials & Correspondence

Indicate the author to whom correspondence and material requests should be addressed.

## Methods

### Specimen collection

Adult *Doliops similis* were collected from Lanyu Island (22°4’44.57”N, 121°33’15.25”E), Taiwan. Dry samples were obtained from fresh ones immersed in 70% ethanol and desiccated at room temperature.

### Optical microscopy

Images of the appearances of adult beetles and scales on black cuticles were taken with a Leica DVM6 digital microscope equipped with the objectives PlanAPO FOV 12.55. Angle-resolved specular spectra (θ_in_ = θ_out_ is measured in the range θ_in_ = 0° to 60°) of the green stripes on the elytra of beetles were collected using an optical goniometer setup: a 5 mm collimated beam was produced by a xenon lamp (HPX2000, Ocean Optics) coupled with an optical fibre (Thorlabs, FC-UV100-2-SR) to a reflective collimator (RC08SMA-F01). The same type of reflective collimator and optical fibre was coupled to a spectrometer (Avantes HS2048) for detection. Optical microscopy on each single scale was performed using a Zeiss Axio Scope optical microscope in Köhler illumination equipped with a 50X SLWD objective (Nikon, NA=0.4) coupled to a spectrometer (Avantes HS2048) via an optical fiber (Thorlabs, FC-UV100-2-SR) (detected spot with diameter of a few *μ*m). Spectra were normalized with a silver mirror (Thorlabs, PF10-03-P01).

### Reflective index and reflection wavelength calculations

Reported values include n=1.64 for amorphous calcium phosphate^46^, n=1.58 for amorphous calcium carbonate^47^, and n=1.41-1.43 for fatty acids in insects^48^, n=1.57-1.75 for proteins^49^; and n=1.52-1.61 for chitin^50^. The upper and lower bound of refractive index of the organic phases in PC (*n*_*o,pC*_) can be estimated:

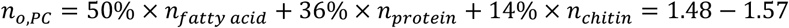

Weighing with minerals (organics/mineral is calculated by relative contribution to O at%), the upper and lower bound of the refractive index of PC (*n*_*pC*_) can be calculated:

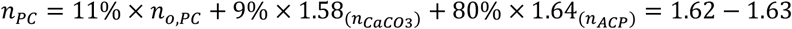

The effective refractive index (*n*_*e f f*_) of the PC and the air spaces is:

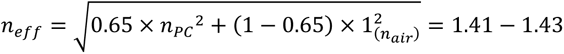

Inserting the average d_111_= 187.9 nm and *n*_*e f f*_ into Bragg-Snell’s law, the predicted reflection peak is at:

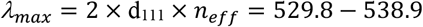

### Focus ion beam scanning electron microscopy (FIB/SEM)

Focus Ion Beam Scanning Electron Microscope (FIB/SEM, Crossbeam 540, Zeiss) was used to collect sequential images with isotropic voxel size of 5 nm. SEM imaging was performed at 2.5 kV acceleration voltage and 700 pA probe current using the ESB electron detector, while slices were generated using the 3 nA FIB probe at a 30 kV accelerating voltage. The acquired dataset was 7160×1680× 540 voxels with a pixel resolution of 5 nm. This dataset was then binned by an isotropic factor of 2 so that the final 3D model had a size of 3580× 840× 270 voxels and 10 nm resolution. Data were segmented by binarization after n2v denoising^51^ and oriented domains were extracted using customized pipeline with an open-source bmmltools (bmmltools, https://bmmltools.readthedocs.io/en/latest/index.html).

### Fourier-transform Infrared Spectroscopy (FTIR)

FTIR spectra of scales were collected using micro-spectrometer (LUMOS II, Bruker) equipped with Mercury Cadmium Telluride (MCT) detector under transmission mode. Scales were taken from the beetle elytra and placed on an IR-transparent KBr window material. To keep the environment stable and lower the noise signals from water vapor or CO_2_, the sample stage is kept in a customized chamber in which N_2_ is continuously purged.

### Small- / Wide-angle x-ray scattering (SAXS/WAXS)

Scales were taken off from the beetles’ elytra and mounted with water on silicon nitride membranes (Norcada Inc., membrane: 2.7 mm× 2.7 mm, 1000 nm thick). SAXS/WAXS 2D diffraction patterns of individual scales^52^ were acquired (experiment SC-5470) at the the ID13 beamline Synchrotron ESRF nanobranch (Grenoble, France). Data with a broad q range from 0.17-41 nm^-1^ were collected with a beamsize of 300-400 nm at an energy of 15 keV with multi-bunch mode. Scan steps are either 300 nm or 500 nm with exposure times varied from 10 ms to 30 ms. 2D patterns were acquired in transmission mode using a Dectris, EIGER X 4M detector.

### X-ray photoelectron spectroscopy (XPS)

XPS was performed with a Thermo Scientific Nexsa spectrometer (Thermo Scientific, UK) using a monochromated Al Kα (*hυ*= 1486.6 eV). Surface charging was mitigated by the use of the charge compensation capability, which utilizes both low energy Ar^+^ ions and low energy electrons. As some components formed in the weevil might be organic in nature, profiling through the scale was achieved by sputtering with the cluster-ion beam (MAGCIS) at a beam energy of 6,000 eV and a cluster size of 1,000 Ar atoms, with spectra recorded using a 100 μm X-ray spot and the snapshot spectral acquisition mode. No additional sample preparation was carried out prior to data acquisition. Data was analysed using AvantageTM V6 (https://mymicroscope.thermofisher.com/public-downloads), where quantitative analysis was performed using the 2p peaks of Ca and P, 2s peak of Mg and 1s peaks of F, N, O and C after fitting with a Shirley background. Three selected etch levels for further chemical analyses were based on the observation of sample surface with the light microscope built in the XPS also where the element profile reaches a plateau.

### Time of Flight Mass Spectrometry (ToF-SIMS)

The ToF-SIMS experiments were performed using a commercial ToF-SIMS mass spectrometer (IONTOF”TOF.SIMS^5^-300”). Both positive and negative ion modes of image acquisition were used with the mass range of m/z= 0-580. The data acquisition software used was SurfaceLab 7.3 (ION-TOF GmbH, Münster, Germany). Experiments were performed and data were analysed by TASCON GmbH (Münster, Germany). The TOF-SIMS chemical characterization of the resulting crater bottom were performed by using Bi_3_^+^ primary ion beam (30 keV). Depth profiles was achieved by sputtering the organic materials with low energy Ar^+^ cluster beam (5,000 eV and a cluster size of 1,000 Ar^+^). Mass resolved secondary ion images (imaging with chemical mapping) were acquired by probing the focused primary ion beam on the surface of interest. For each pixel addressed, a complete spectrum is recorded. Field of view of 150×150 µm with lateral resolution of 500 nm and nominal resolution of 100 nm was achieved by rastering of the primary ion beam. The m/z profiles provide ToF-SIMS depth profiling and imaging were conducted, collecting positively and negatively charged ions within mass ranges from m/z= 0 to m/z=580. The per pixel mass spectra are normalized with the sum of mass counts at that pixel

### Transmission electron microscopy-energy-dispersive X-ray spectroscopy/ electron energy loss spectroscopy (TEM-EDS/EELS)

Fragments of cuticles with scales were dehydrated through a graded ethanol series (70–100%) followed by infiltration in Embed812 epoxy resin (Electron Microscopy Sciences) for 5 days. The resin blocks containing the specimens were then hardened at 70 °C for 24 hours. Ultrathin sections (thickness of ca. 70 nm) of the embedded specimens were prepared using an EM UC7 ultramicrotome equipped with a 45° diamond knife (Diatomes). The sections were collected from an ethylene glycol-filled knife boat and transferred to lacey carbon-supported Cu grids^53^. The grids were vacuum-dried at room temperature before TEM observations. Transmission electron microscopy have been conducted using a JEOL JEM F200 operated at 200 kV acceleration voltage equipped with a GATAN OneView CMOS camera for fast imaging. Local EDS analysis was performed using a dual 100 mm^2^ window-less silicon drift detector. Electron Energy Loss Spectroscopy (EELS) have been performed using a GATAN GIF Continuum ER spectrometer, a convergence semi-angle of 10 mrad and a collection semi-angle of 4.2 mrad.

