## Supplement Information for "Longhorn Beetles Form Structural Colour Using Calcium Phosphate Biominerals"

### Methods

#### Focus ion beam scanning electron microscopy (FIB/SEM)

FIB-SEM sequential images with isotropic voxel size of 10 nm were collected. SEM imaging was performed at 2.5 kV acceleration voltage and 700 pA probe current using ESB electron detector, while slices were generated using the 3 nA FIB probe at a 30 kV accelerating voltage.

#### Scanning electron microscopy energy dispersive X-ray spectrometry (FIB/SEM-EDS)

Scales were taken from different beetles and put separately on the carbon tape on a SEM stub. Each scale was trenched with FIB ion gun along the cross-section to expose the inner photonic crystals. Data was acquired using a Crossbeam 550 (Zeiss, Jena, Germany) equipped with a Ultim Extreme detector (Oxford instruments, Oxfordshire, UK). Accelerated voltages of 15 keV were applied to the samples. Acquisition was performed for 20 minutes with process time of 4. Results were analyzed using AZtech software (Oxford instruments, Oxfordshire, UK). The weight percentage of detected elements in each sample were presented as pie chart.

#### X-ray photoelectron spectroscopy (XPS) of pure standards

Pure standards of hydroxyapatite (HAP), α-tricalcium phosphate (α-TCP), calcium carbonate minerals $CaCO_3$ were purchased from Sigma Aldrich. Amorphous calcium phosphate (ACP) was synthesized followed the reported method[1]. The insect fatty acid reference was extracted from the surface of dustwings. Fibroin proteins were extracted and purified from silk cocoon. Chitin reference was extracted and purified from crabs. XPS was performed with a Thermo Scientific Nexsa spectrometer (Thermo Scientific, UK) using a monochromated Al Kα ($h\upsilon$ = 1486.6 eV). Spectra of standards were recorded using a 400 μm X-ray spot and the scanned spectral acquisition mode.

#### Scanning transmission electron microscopy- energy-dispersive X-ray spectroscopy quantification (STEM-EDS)

Atomic percentage of elements per pixel were calculated using GATAN software applying the Cliff-Lorimer ratio equation.

#### Small- / Wide- angle x-ray scattering (SAXS/WAXS)

SAXS/WAXS 2D diffraction patterns of individual scales from different rotation angles ($0^{\circ}$ -$30^{\circ}$) were acquired in the experiment SC-5328) on the microbranch of ID13 beamline Synchrotron ESRF microbranch (Grenoble, France). Data with a broad q range from 0.17- 44 $nm^{-1}$ were collected with a beamsize of 2.5× 2.5 $\mu m^2$ at an energy of 13 keV with multi-bunch mode. Scan steps varied between 1- 2 μm with exposure times from 10 ms to 25 ms. 2D patterns were acquired in transmission mode using a Dectris, EIGER X 4M detector. Scales were taken off from the beetles' elytra and mounted with water on silicon nitride membranes (Silson Ltd., membrane: 10 mm × 10 mm, 200 μm thick) and fixed on a rotation stage.

### 1. Structural Heterogeneity in the Photonic Crystals

#### 1.1 Crystal Symmetry

The green pattern of *D. similis* is covered with stacked, micro-sized scales. Each scale is connected to the cuticle, 50 μm deep, by a narrow stalk 1 μm in diameter, and protrudes from the cuticle toward the posterior.

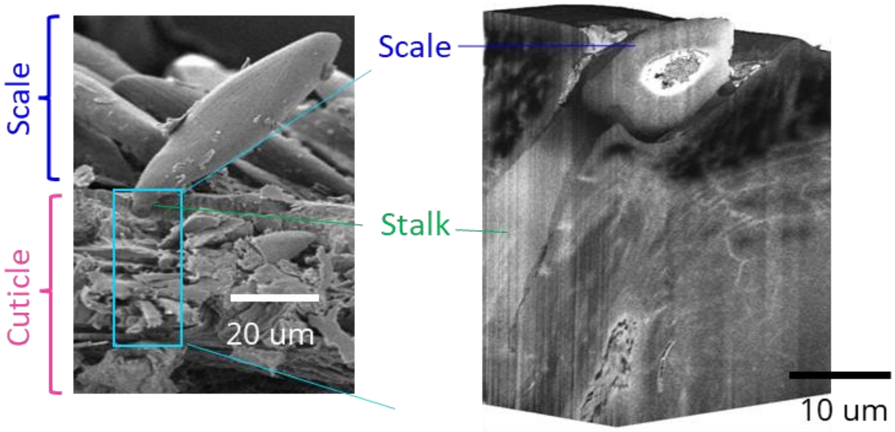


**Fig. S1:** Left: 2D SEM image of a broken beetle shell reveals the connection stalk of a scale. Right: 3D volume image of a scale stalk in the cuticle.

Face-centered cubic (FCC) structures differ from hexagonal close packed (HCP) structures only in stacking order: both structures have close-packed atomic planes along [111] and [001], respectively. When stacking one of these layers on top of another, the units in successive layers are not directly on top of one another. The first two layers are identical for HCP and FCC, and labelled AB. If the third layer is placed such that its units are directly above those of the first layer, the stacking will be ABA — this is the HCP structure. In the FCC structure, the third layer of units is arranged differently than the first and second layers. Leading to ABCABC stacking along the [111] direction (**Fig. S2** upper panel).

In the **Fig.S2** lower panel, three continuous layers of (111) are extracted from different regions in the same domain, and every two neighboring layers are stacked with colour code red and green. Black space refers to empty space without particles. At the site 1 of the domain 1, the stack of layer 1 and 2 (left image) is complemented with the stack of layer 2 and 3 (right image) which indicates a FCC stacking that atoms in layer 3 fill in different voids from layer 1. While at the site 2 of the domain 1, the stack of layer 1 and layer 2 (left image) is similar to the stack of layer 2 and layer 3 (right image) which indicates a HCP stacking.

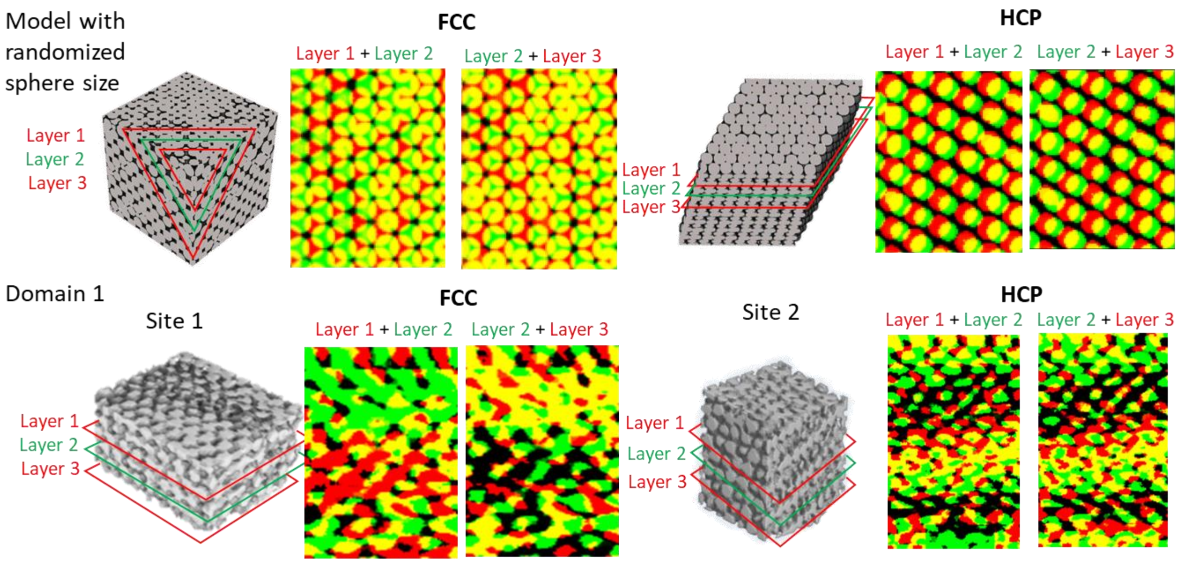


**Fig. S2:** Mixture FCC and HCP stacking locally in the same domain. (surface normal of [111], and [001] respectively). Upper panel shows the stacking of neighboring layers (coloured with red and green, respectively. Overlap between the layers is denoted in yellow) along [111] in FCC and [001] HCP models with randomized sphere size. Lower panel: Overlay patterns of two sites of the PC are presented. Site 1 in domain 1 is more similar to FCC, whereas site 2 in the same domain shares higher similarity with HCP.

#### 1.2 Structural Parameters

After successfully segmenting domains in the photonic crystals with the similar structural features, we further calculate structural parameters of each domain. The calculated filling fraction (FF), d-space of (111) crystal planes, lattice constant ($a$), and the diameter of nanospheres ($\emptyset$) of each domain is listed in Table S1.

Diffraction from (111) is the most intense peak in the integration spectrum of FFT. We measured the q-value of the (111) and its standard deviation δq then calculated the d-space of (111) with deviation $\sigma_{d_{111}}$in each domain. Based on geometry of the FCC lattice, we derive the lattice constant $a \pm \delta_a$ from the $d_{111} \pm \sigma_{d_{111}}$

$$d_{111} = \frac{1}{q}$$

$$\sigma_{d_{111}} = \frac{\delta q}{q^2}$$

$$a = \sqrt{3}d_{111}$$

$$\delta_a = \sqrt{3}\sigma_{d_{111}}$$

Filling fraction (FF) is defined as the volumetric ratio of nanospheres (with diameter $\emptyset$) to the repeated FCC lattice, calculated in the Dragonfly software (Dragonfly 2022.2.0.1409, Object Research Systems (ORS) Inc., Montreal, Canada, 2022; software available at https://www.theobjects.com/dragonfly). by dividing the total voxels of nanospheres by the voxels of the domain:

$$FF = \frac{4 \times \frac{4}{3}\pi \times (\frac{\emptyset}{2})^3}{a^3}$$

Deviation of filling fraction δF is calculated with different brightness threshold in the Dragonfly software:

$$\delta F = FF_{up} - FF_{low}$$

Nanosphere diameter, thus, can be derived from the measured FF (non-close packed):

$$\emptyset = \sqrt[3]{\frac{3 \times FF}{2\pi}}a$$

$$\delta\emptyset = \sqrt{\left(\frac{a}{3} \times \left(\frac{2\pi}{3 \times FF}\right)^{\frac{2}{3}} \times \delta F\right)^2 + \left(\left(\frac{3 \times FF}{2\pi}\right)^{\frac{1}{3}} \times \delta_a\right)^2}$$

**Table S1. Structural parameter of each domain in the photonic crystals in a scale.**

| | FF (measured) | δF (measured) | q-value (measured) | $d_{111}$ (nm) | $a$ (nm) | ∅ (nm) |
|---|---|---|---|---|---|---|
| Domain1 | 0.55 | 0.1 | 0.0055±0.0022 | 181.8±72.7 | 314.9±125.9 | 201.7±84.6 |
| Domain2 | 0.67 | 0.2 | 0.0055±0.0020 | 181.8±66.1 | 314.9±114.5 | 215.4±90.3 |
| Domain3 | 0.66 | 0.2 | 0.0056±0.0019 | 178.6±60.6 | 309.9±104.9 | 210.9±84.2 |
| Domain4 | 0.63 | 0.2 | 0.0053±0.0019 | 188.7±67.6 | 326.8±117.2 | 219.5±92.3 |
| Domain5 | 0.64 | 0.2 | 0.0052± 0.0016 | 192.3±59.2 | 333.1±102.5 | 224.3±84.6 |
| Domain6 | 0.63 | 0.2 | 0.0049± 0.0015 | 204.1±62.5 | 353.5±108.2 | 236.9±89.5 |

Higher disorder is observed in the domain 6 at the place closer to the end of the tail of a scale (**Fig. S3**a, white area). Viewing from the end tail of domain 6 (**Fig. S3**b) shows varied nanosphere sizes at the tail of the scale.

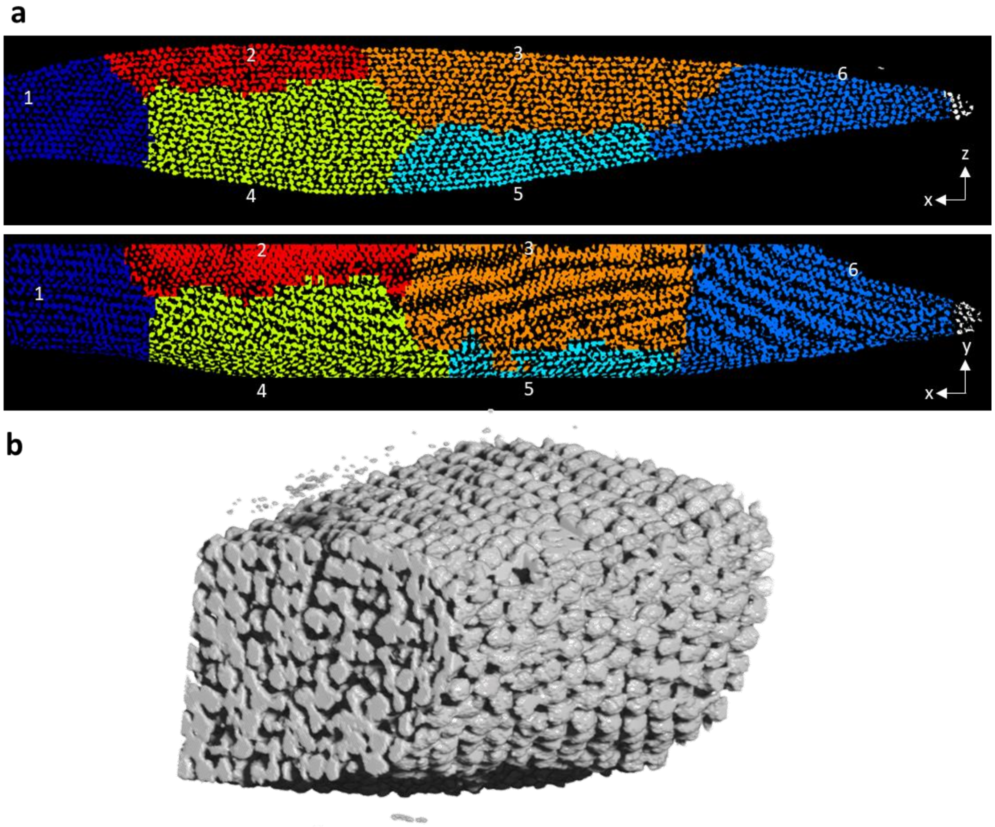


**Fig. S3:** Increased disorder in domain 6 at the tail of the scale. a, Frontal plane and sagittal plane views of the segmented photonic crystal in an intact scale (each domain is colored differently). b, a 3D model of domain 6.

### 2. Compositions analysis

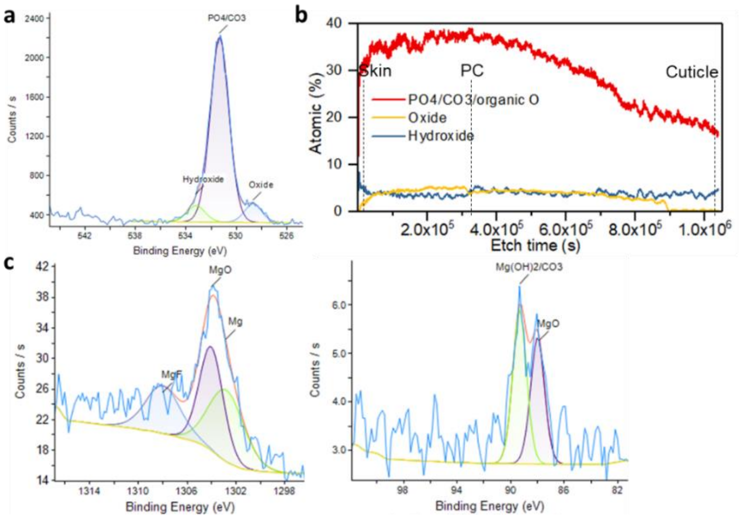

**Fig. S4:** a, Deconvolution of O 1s spectra with hydroxide, oxide, and other oxygen species and b, the calculated depth profile of oxygen contributed from different species. c, Peak fitting Mg 1s, Mg 2s spectra detected in the photonic crystal region.

**Table S2. Ca/P and O/P ratios in the skin, photonic crystals (PC), cuticles and pure standards of calcium phosphate minerals (HAP, α-TCP, ACP), calcium carbonate minerals $CaCO_3$.**

| | Ca2p1 | Ca2p3 | P2p | Ca /P | O/P |
|---|---|---|---|---|---|
| Skin | 350.8 | 347.2 | 133.5 | 1.3 | 4.1 |
| PC | 350.9 | 347.3 | 133.5 | 1.5 | 4.0 |
| Cuticle | 351.1 | 347.4 | 133.5 | 1.9 | 5.5 |
| HAP | 351.3 | 347.8 | 133.5 | 1.4 (1.67) | 4.4 |
| α-TCP | 351.3 | 347.8 | 133.2 | 1.2 (1.5) | 3.7 |
| ACP | 350.9 | 347.3 | 133.4 | 1.0 (1.5) | 4.4 |

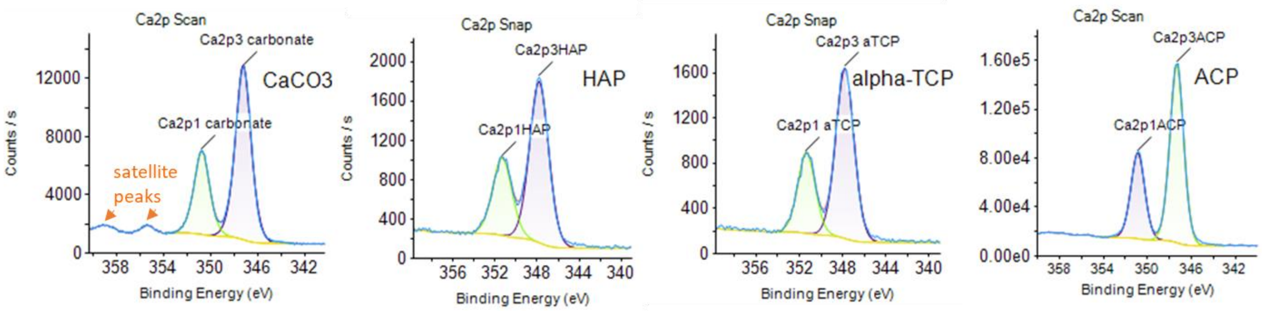


**Fig. S5:** XPS Ca 2p of pure mineral standards. The table listed the peak fitting energy of Ca 2p3, Ca 2p1, P 2p in skin, photonic crystals, cuticles, and standards.

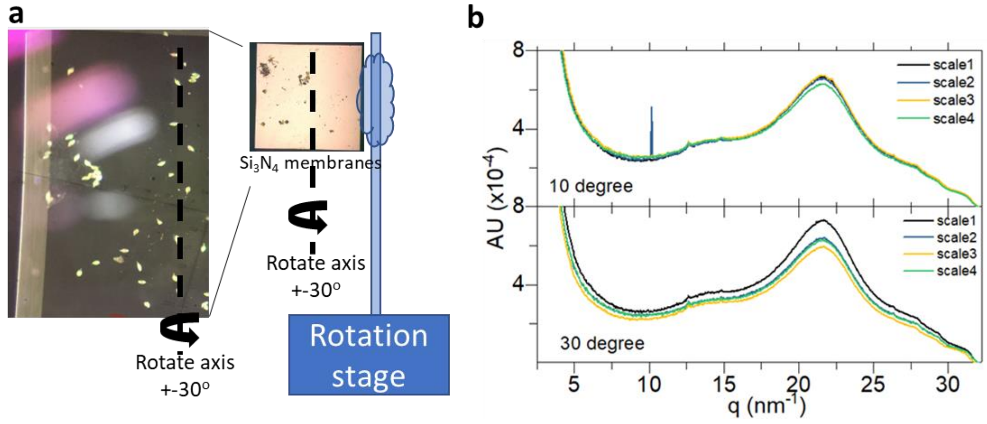


**Fig. S6:** SAXS/WAXS from the *D.similis* scale with different angle of incident X-ray on individual scale. a, Experimental setup for measuring the diffraction signals with varied angles of ±30°. b, Integration XRD signals of 4 individual scales with different rotation angles[2].

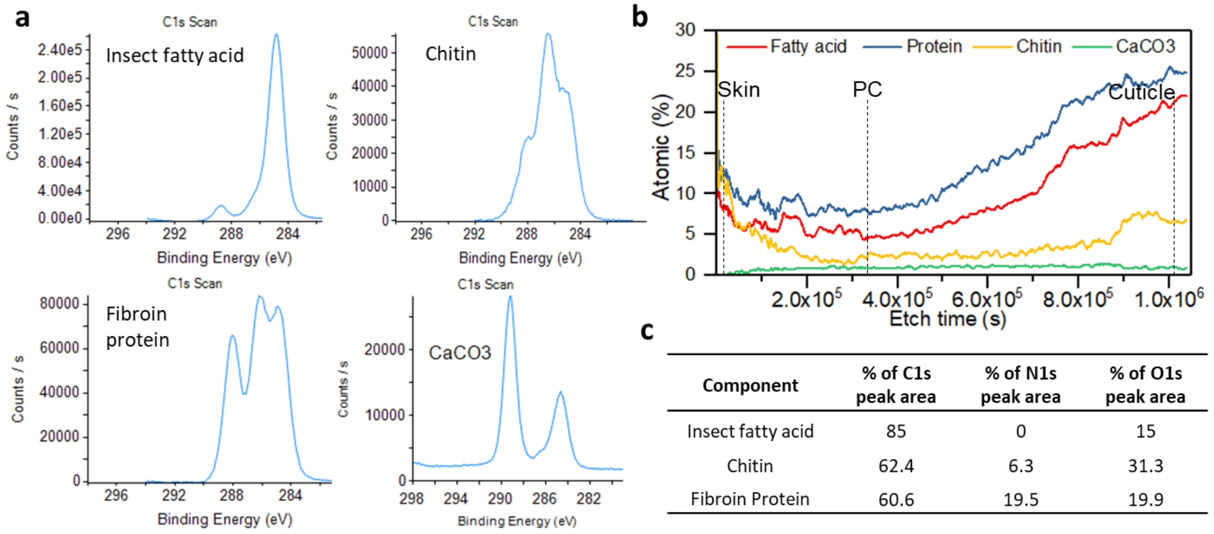


| Component | % of C1s peak area | % of N1s peak area | % of O1s peak area |
|---|---|---|---|
| Insect fatty acid | 85 | 0 | 15 |
| Chitin | 62.4 | 6.3 | 31.3 |
| Fibroin Protein | 60.6 | 19.5 | 19.9 |

**Fig. S7:** a, XPS C 1s spectrum of pure standards: fatty acids (collect from insect wax), protein (purified fibroin proteins), chitin (purified a-chitin from crab), calcium carbonate. b, Depth profile of components derived from multivariate analysis. c, Amounts of C, N, O in the multivariate references quantified by integrating XPS peak area.

**Table S3. Theoretical ratio of C, N, O, in PC derived from the % of C 1s and elemental contents in PC (Fig. 2c).**

| Component | % of C1s peak area | C content at% (Theoretical) | N content at% (Theoretical) | O content at% (Theoretical) |
|---|---|---|---|---|
| Insect Fatty Acid | 47.0 | 7.8 | 0 | 1.4 |
| Alpha Chitin | 12.5 | 2.1 | 0.2 | 1.1 |
| Fibroin Protein | 34.1 | 5.6 | 1.8 | 1.8 |
| $CaCO_3$ | 6.4 | 1.1 | 0 | 3.3 |

To examine the composition homogeneity, we performed RPCA on the per pixel normalized TEM-EELS spectrum with energy loss from 220 eV to 500 eV, covering the C K edge, Ca K edge, and N kedge after removing the Lacy-C support layer. The most significant three decomposition factors and their loadings were shown in Fig. S8. The most intense pixels in the three decomposition factors were selected and converted to three individual binary masks for extracting the average EELS and EDS spectra.

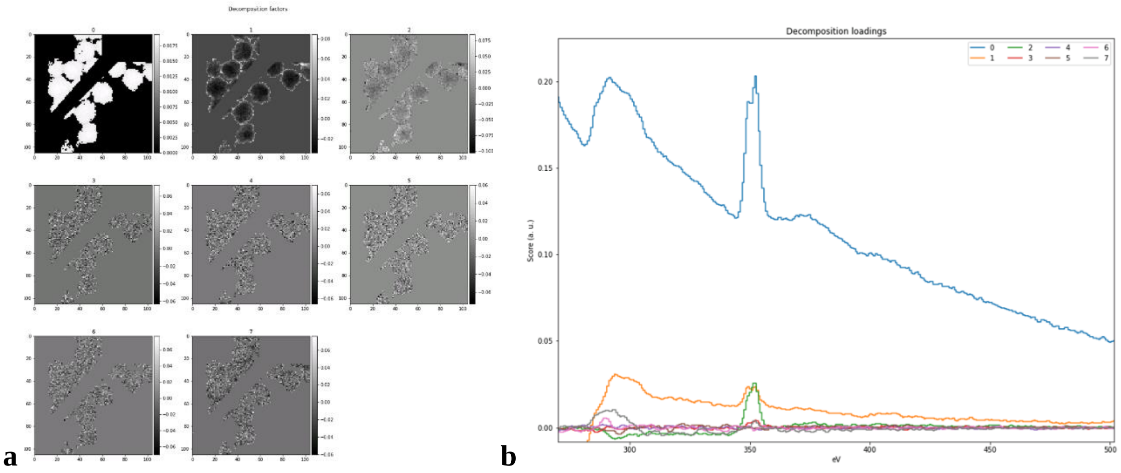


**SI Fig. S8** a, RPCA decompose factors and b, the loadings of normalized EELS hyperspectra. The area of/overlapped with LacyC was masked out before decomposition analysis. Numbers of factor are set to 8. Decomposition factor 0, 1were selected to generate binary masks by thresholding the brightest pixels in each factor, which were used for further composition analyses.

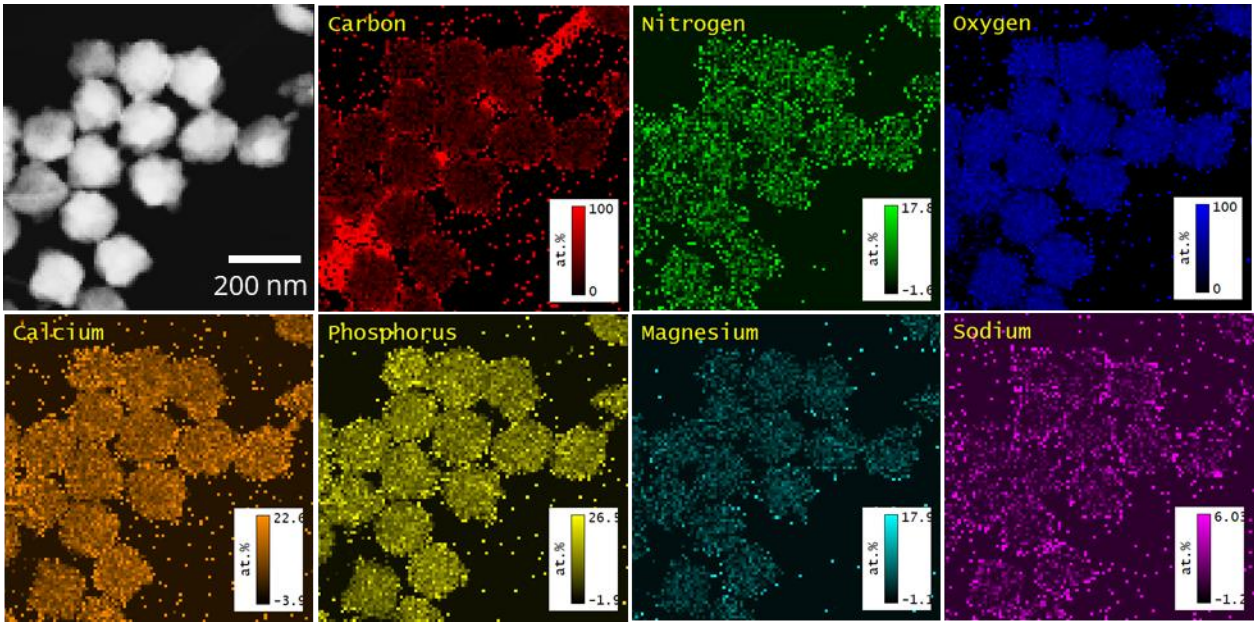


**SI Fig.** S9 STEM-HAADF (high angle annular dark field) image and the TEM-EDS element atomic percentage mapping.

STEM-HAADF gives a contrast resulted from differences in atomic number and local thickness. Higher atomic number and thicker places are brighter. The LacyC support network is seen with the highest C at%. The outer surface of each nanosphere is with higher C at% and N at% at some places. The distribution of Na also shows higher percentage at the outer shell.

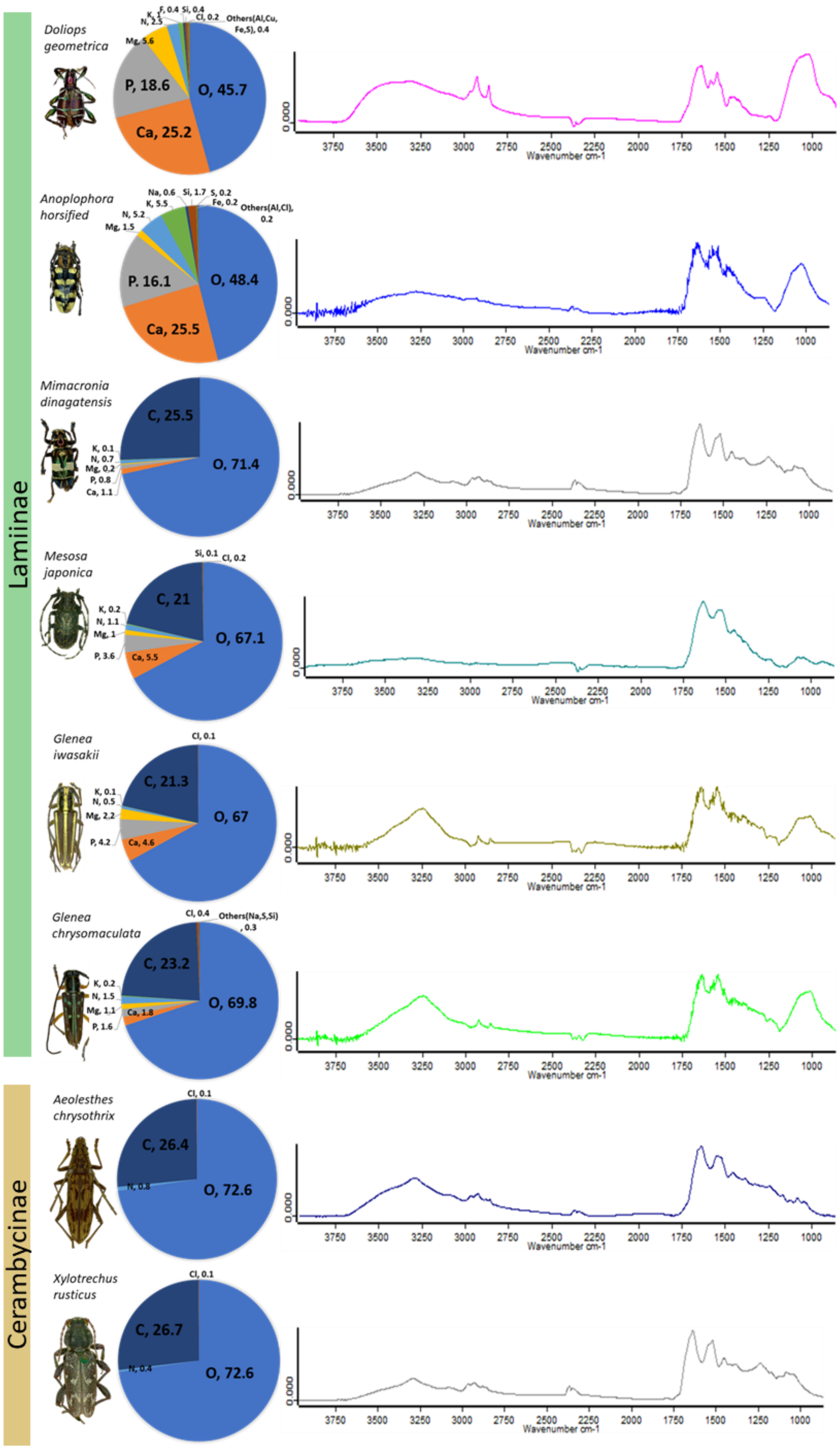

Laminae
Cerambycinae
Doliops geometrica
O, 45.7
Ca, 25.2
P, 18.6
Mg, 5.6
N, 2.5
F, 0.4
Si, 0.4
Cl, 0.2
Others(Al,Cu, Fe,S), 0.4
Anoplophora horsified
O, 48.4
Ca, 25.5
P. 16.1
Mg, 1.5
N, 5.2
K, 5.5
Na, 0.6
Si, 1.7
S, 0.2
Fe, 0.2
Others(Al,Cl), 0.2
Mimacronia dinagatensis
O, 71.4
C, 25.5
K, 0.1
N, 0.7
Mg, 0.2
P, 0.8
Ca, 1.1
Mesosa japonica
O, 67.1
C, 21
Si, 0.1
Cl, 0.2
K, 0.2
N, 1.1
Mg, 1
P, 3.6
Ca, 5.5
Glenea iwasakii
O, 67
C, 21.3
Cl, 0.1
K, 0.1
N, 0.5
Mg, 2.2
P, 4.2
Ca, 4.6
Glenea chrysomaculata
O, 69.8
C, 23.2
Cl, 0.4
Others(Na,S,Si), 0.3
K, 0.2
N, 1.5
Mg, 1.1
P, 1.6
Ca, 1.8
Aeolesthes chrysothrix
O, 72.6
C, 26.4
Cl, 0.1
N, 0.8
Xylotrechus rusticus
O, 72.6
C, 26.7
Cl, 0.1
N, 0.4
0.000
3750 3500 3250 3000 2750 2500 2250 2000 1750 1500 1250 1000
Wavenumber cm-1

**SI Fig. S10** Element weight percentage calculated from SEM-EDS spectra and FTIR of scales from differently coloured Lamiinae longhorn beetles.
